# Complement receptor C5aR1 signaling in sensory neuron-associated macrophages drives neuropathic pain

**DOI:** 10.1101/2022.07.01.498487

**Authors:** Andreza U. Quadros, Alexandre G. M. Maganin, Conceição E. A. Silva, Samara Damasceno, Maria C. M. Cavallini, Marcela Davoli-Ferreira, Alexandre H. P. Lopes, Devi R. Sagar, Laura Brandolini, Sang Hoon Lee, Jose C. Alves-Filho, Fernando Q. Cunha, Temugin Berta, Jörg Köhl, Marcello Allegretti, Victoria Chapman, Thiago M. Cunha

**Affiliations:** Center for Research in Inflammatory Diseases (CRID), Department of Pharmacology, Ribeirao Preto Medical School, University of Sao Paulo, Avenida Bandeirantes, 3900, 14049-900, Ribeirao Preto, SP, Brazil; Graduate Program in Pharmacology Ribeirao Preto Medical School, University of Sao Paulo, Avenida Bandeirantes, 3900, 14049-900, Ribeirao Preto, SP, Brazil; Graduate Program in Basic and Applied Immunology Ribeirao Preto Medical School, University of Sao Paulo, Avenida Bandeirantes, 3900, 14049-900, Ribeirao Preto, SP, Brazil; Pain Centre Versus Arthritis, School of Life Sciences, University of Nottingham, Clinical Sciences Building City Hospital, Nottingham, NG5 1PB, United Kingdom; R&D Department, Dompé Farmaceutici s.p.a., via Campo di Pile, 67100 L’Aquila, Italy; Pain Research Center, Department of Anesthesiology, University of Cincinnati College of Medicine, Cincinnati, Ohio, 45267, USA; Institute for Systemic Inflammation Research, University of Lübeck, Ratzebuger Allee 160 23562 Lübeck, Germany; Division of Immunobiology, Cincinnati Children’s Hospital Medical Center and University of Cincinnati College of Medicine, Cincinnati, OH, 45229, USA

**Keywords:** neuropathic pain, complement system, macrophages, C5aR1, IL-1b

## Abstract

Neuroimmune interactions across the pain pathway play a predominant role in the development of neuropathic pain. Previous reports demonstrated that complement driven effector systems including the C5a/C5aR1 axis contribute to these neuro-immune mechanisms. However, the cellular and molecular mechanisms underlying C5a/C5aR1 signaling-mediated neuropathic pain development remain ill-identified. Here we show that neuropathic pain following peripheral nerve injury was attenuated in C5aR1-deficient male and female mice as well as in wild type mice treated with a selective allosteric C5aR1 antagonist. Using two complementary cell-specific C5aR1 knockout mouse strains, we identified C5a/C5aR1 driven-activation of sensory neuron-associated macrophages (sNAMs) located in the sensory ganglia as the key site of peripheral nerve injury-induced neuropathic pain, whereas activation of macrophages of the local of peripheral nerve injury was not involved. Mechanistically, we uncovered IL-1b the main mediator of pain hypersensitivity in response to C5aR1 signaling in sNAMs. Our findings highlight a crucial role of C5a/C5aR1 axis activation in sNAMs for the development of neuropathic pain and identify this pathway as a promising novel target for neuropathic pain therapy.

## Introduction

Neuropathic pain is a subtype of chronic pain caused by injury or disease affecting the somatosensory system^1,2^. Although knowledge of the underlying pathophysiological mechanisms of neuropathic pain has increased in the last decades, the current treatment and management of neuropathic pain is still a challenge^2,3^. The discovery of fundamental and specific mechanisms that mediate the induction and maintenance of neuropathic pain is essential for the development of a novel pipeline of pharmacological treatments. Mounting evidence supports the crucial role of neuroimmune glial in nociceptive processing and the development and maintenance of neuropathic pain, especially those caused by peripheral nerve injury^4–10^.

Following a peripheral nerve injury, there is an immediate inflammatory/immune response at the injury site, which is characterized by the earlier recruitment of neutrophils followed by the macrophages/monocytes and CD4+ T cells infiltration that can play a role in the development of neuropathic pain^4,9–11, 67^.The recruitment of these cells into the site of nerve injury is governed by the release of several inflammatory mediators, including cytokines and chemokines^6,9,12,13^. Together, these immune cells and inflammatory mediators likely contribute to the sensitization of primary sensory neurons and consequently to the manifestation of neuropathic pain. Peripheral nerve injury also triggers distal neuroimmune glial interactions in the sensory ganglia [dorsal root ganglions (DRGs) and trigeminal ganglion, (TG)] and in the dorsal horn of the spinal cord. In the sensory ganglia, there is an increased activation/ proliferation of macrophages and satellite glial cells, that in turn produce pro-nociceptive mediators, such as TNF and IL-1β^4,10,14–16^.

Neuro-immune glial interactions driving neuropathic pain are multifactorial, and inflammatory mediators beyond cytokines also play an important role^6,9,12^. Molecules belonging to the complement system, especially C5a, have been implicated in various pain conditions^17–19^. C5a is one of the most important mediators of the complement cascade, which possesses several pro-inflammatory actions^20–23^. C5a is generated in response to activation of all complement pathways^24,25^ and confers it effector functions mainly via the GPCR C5a receptor type 1 (C5aR1), also called CD88^26,27^. C5aR1 was initially identified in neutrophils, monocytes/macrophages, and mast cells^28–31^. C5a/C5aR1 signaling has been implicated in the genesis of acute (post-operative)^32–34^, inflammatory^35^, and neuropathic pain conditions^36,37^. Although previous work has implicated C5a/C5aR1 signaling in the development of neuropathic pain^19,36^, the underlying mechanisms are incompletely understood^19^. Herein, we confirmed that C5a/C5aR1 signaling is required for the development of neuropathic pain after peripheral nerve injury. In particular, we identified a previously undetermined role of this signaling in sensory neuron-associated macrophages (sNAMs) located in the sensory ganglia. We also found that C5a/C5aR1 signaling in sNAMs triggered the production of the pro-nociceptive mediator IL-1β. Furthermore, inhibition of C5a/C5aR1 signaling significantly reduced the development of neuropathic pain in male and female mice after peripheral nerve injury.

## Results

### The C5aR1 signaling is involved in the development of neuropathic pain

Baseline nociceptive behaviors were assessed in WT and *C5ar1*^-/-^ male mice after application of different stimuli into the hind paw. No significant differences were detected in *C5ar1*^-/-^ compared to WT mice following exposure to mechanical (von Frey filament and electronic) or thermal (cold; acetone test and heat; Hargreaves and hot plate tests) nociceptive stimuli (**Fig. 1a**). In addition, motor performance (rotarod) was comparable between WT and *C5ar1*^-/-^ mice (**Fig. 1a)**. Next, we investigated whether C5aR1 participated in the development of nerve injury-induced neuropathic pain caused by partial ligature in the common branch of the sciatic nerve (PSNL model). The injured WT mice (WT PSNL) exhibited both mechanical and cold pain hypersensitivity in this model compared to sham-operated WT mice (WT Sham-male mice) (**Fig. 1b**). In contrast, *C5ar1*^-/-^ mice with PSNL (*C5ar1*^-/-^ PSNL-male mice) had reduced levels of mechanical and cold pain hypersensitivity after nerve injury compared to WT PSNL mice (**Fig. 1b**). To evaluate potential gender differences^16,53^, studies were also undertaken in female mice. Similar to male mice, mechanical and cold pain hypersensitivity induced by PSNL was reduced in *C5ar1*^-/-^ female mice (**Supplementary Fig. 1**). Given that we did not observe gender differences, all subsequent experiments were conducted in male mice.

**Figure 1.**
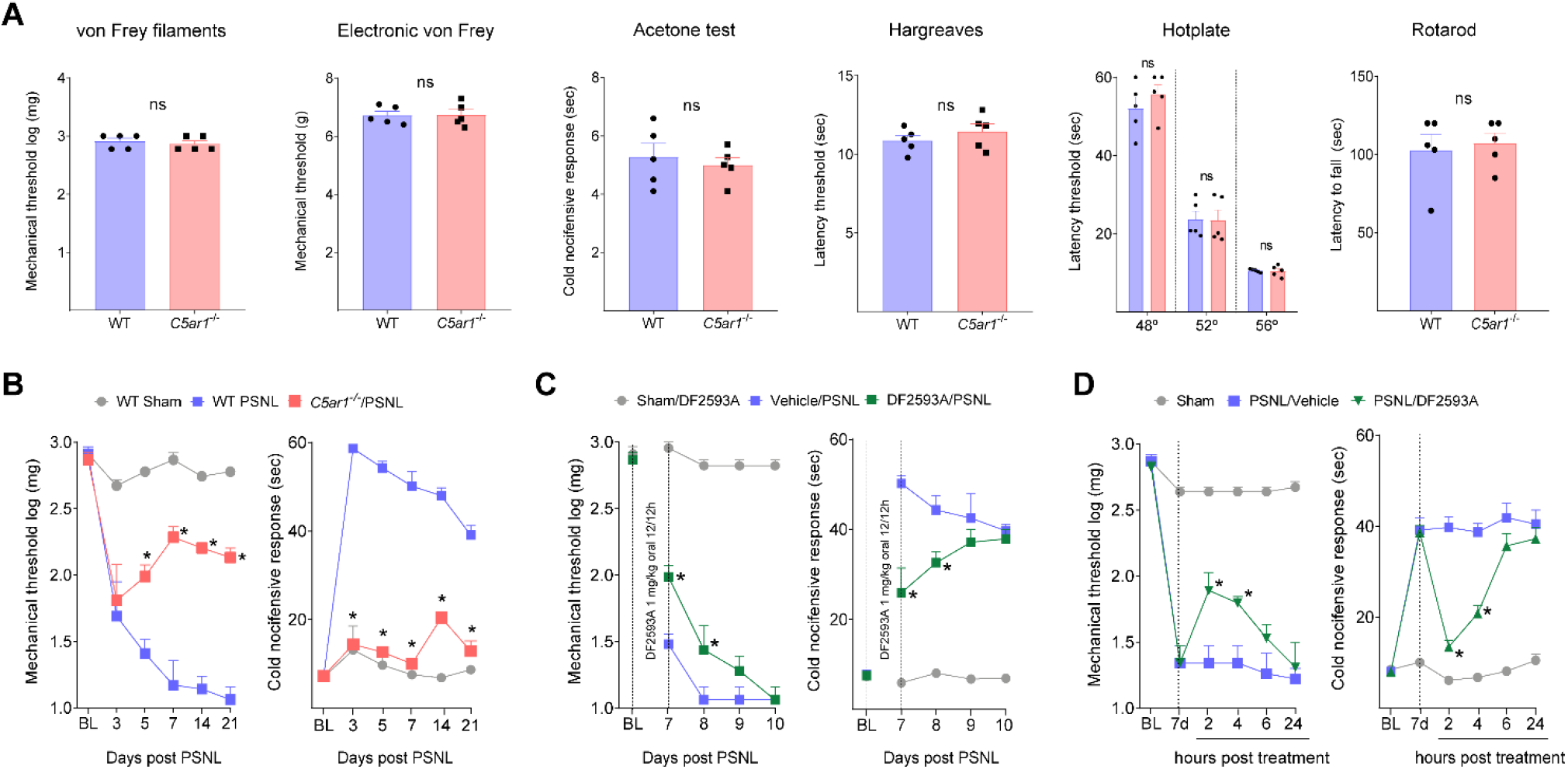
C5aR1 is not involved in nociceptive pain but drives the development of neuropathic pain. **(A)** Physiological nociceptive responses of Balb/C WT or *C5ar1*^*-/-*^ mice to mechanical (von Frey filament and electronic von Frey, *n*=5), and thermal stimulation (Acetone test, Hargreaves and Hotplate, *n*=5) or motor coordination (Rotarod test, *n*=5). Data are represented as the mean ± s.e.m. and analyzed by non-paired t test - ns, non-significant. (**B)** Development of mechanical and cold allodynia in Balb/C WT or *C5ar1*^*-/-*^ mice, from 3 up to 21 days after PSNL surgery. (**C)** Mechanical and cold allodynia in Balb/C WT mice, treated orally twice daily with 1 mg/kg of DF2593A from the day of surgery until the 7^th^ day post PSNL. Nociceptive behaviors were analyzed from 7 up to 10 days after surgery (*n*=6). (**C)** Balb/C WT animals treated in the 7th day post-surgery with a single dose of DF2593A (orally, 1 mg/kg) and mechanical and cold allodynia were analyzed (*n*=6). Data are represented as the mean ± s.e.m. and analyzed by non-paired t test - ns (**A**) non-significant; or by repeated measures Two-Way ANOVA with post hoc Bonferroni (**B-D**). \**P*<0.05 compared to WT/PSNL or vehicle/PSNL group.

Our data suggested that C5aR1 signaling might be a promising target to prevent the development of neuropathic pain, Thus, we tested whether pharmacological inhibition of C5aR1 signaling prevented the development of neuropathic pain. Firstly, systemic treatment (orally, twice a day) of WT mice with a selective allosteric inhibitor of C5aR1 signaling, DF2593A^37^, starting at the day of nerve injury up to 7 days post-PSNL, reduced mechanical and cold pain hypersensitivity induced by PSNL (**Fig. 1c)**. When the treatment was interrupted, at day 7 post-injury, mechanical and cold pain hypersensitivity started to develop in DF2593A-treated animals and reached the same levels of vehicle-treated mice 10 days after PSNL (**Fig. 1c)**. Notably, the post-treatment (therapeutic schedule, single treatment) of mice with DF2593A, at day 7 after PSNL surgery, transiently reduced mechanical and cold pain hypersensitivity compared to vehicle-treated mice (**Fig. 1d)**. Collectively, these data suggest that C5aR1 signaling contributes to the development of neuropathic pain caused by peripheral nerve injury, but is not involved in the acute physiological nociception

### C5aR1 signaling in the site of nerve injury has no major role in neuropathic pain development

In the next part of this study, we sought to investigate the mechanisms by which C5aR1 signaling induces the development of neuropathic pain in response to peripheral nerve injury. Initially, we evaluated a potential role of C5aR1 signaling at the site of nerve injury. The gene expression of *C5ar1* was increased at the site of nerve injury, in a time-dependent manner, peaking at 3 days PSNL induction (**Fig. 2a**). Supporting these results, immunofluorescence analysis revealed an increased number of C5aR1-expressing cells in the local area of peripheral nerve injury (**Fig. 2b-c**). Confirming the specificity of the anti-C5aR1 antibody, no staining was observed in nerve samples from *C5ar1*^-/-^ mice (**Supplementary Fig. 2**).

**Figure 2.**
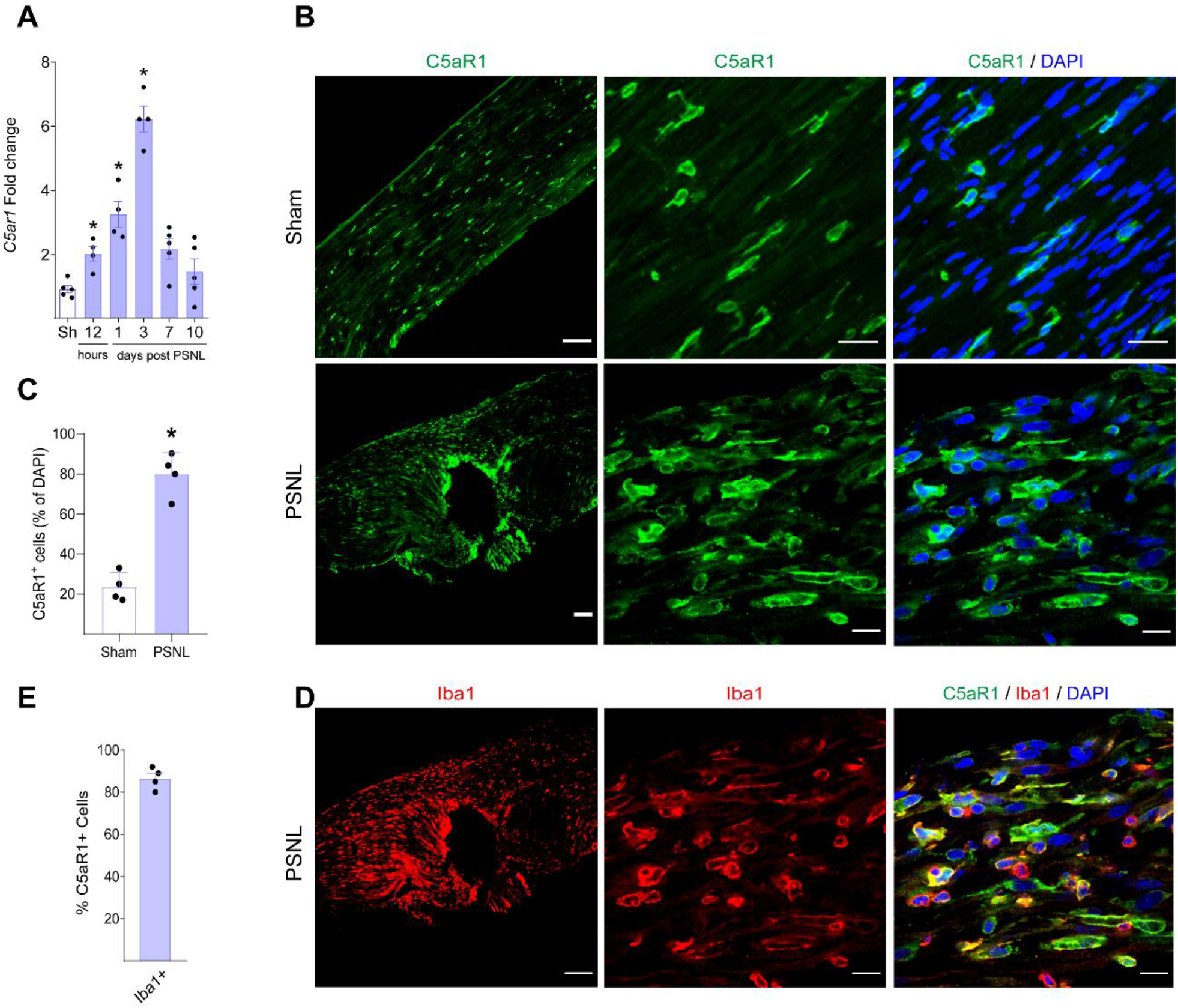
C5aR1 expression is increased in the sciatic nerve after PSNL. **(A)** Sciatic nerve from naive or PSNL mice was harvested and the expression of *C5ar1* gene at indicated time points (qRT-PCR). Data are represented as the mean ± s.e.m., *n*=4-5. \**P*<0.05, related to sham, analyzed by One-Way ANOVA with post hoc Bonferroni. (**B and D)** Representative images and (**C and E)** quantification showing immunoreactivity for C5aR1 (green color) double/triple labeled with DAPI (cell nuclei; blue) and Iba-1 (red color) in the sciatic nerve harvested form Sham or PSNL mice (3 day after surgery), bar scale 30 µm or 100 μm. Data are represented as the mean ± s.e.m., *n*=4. \**P*<0.05 related to sham, analyzed by non-paired t test.

C5aR1 expression has been detected in several cell subtypes, including myeloid cells such as macrophages and neutrophils^28–30^. In order to analyze the cell subtype expressing C5aR1 at the site of peripheral nerve injury, confocal analysis was performed. Double immunostaining revealed that C5aR1 is mainly expressed in cells expressing the canonical marker of macrophages, Iba-1 (86.5%; **Fig. 2d-e**), indicating monocytes/macrophages as the major cellular source of C5aR1 signaling in the local of nerve injury. Supporting these data, C5aR1-GFP mice also revealed the expression C5aR1 in Iba-1 positive cells (**Supplementary Fig. 3a**). On the other hand, C5aR1 expression in the site of nerve injury was detected in few Ly6G+ cells, indicating low expression in infiltrating neutrophils (**Supplementary Fig. 3b**).

Immediately after a peripheral nerve injury, we previously observed a massive infiltration of neutrophils followed by the infiltration of monocytes/macrophages around the injury^54–57^. In support of these findings. flow cytometric analysis of nerve injured tissue showed that there is a significant increase in the number of leukocytes (CD45+ cells; **Supplementary Fig 4a**). The leukocytes were composed mainly of neutrophils (CD11b+ Ly6G+ cells) and monocytes/macrophages (CD11b+ Ly6G-cells) (**Supplementary Fig. 4a**). Since C5a/C5aR1 signaling is involved in the recruitment of immune cells to the site of inflammation^30,34^, we investigated whether this signaling is involved in the recruitment of leukocytes to the site of peripheral nerve injury. Surprisingly, we observed no difference in leukocyte (CD45+ cell) infiltration in the local of peripheral nerve injury of *C5ar1*^-/-^ mice compared to WT mice in all evaluated time points (**Supplementary Fig. 4a and b**). No significant differences between *C5ar1*^-/-^ and WT mice were detected in the infiltration of both neutrophils (CD11b+ Ly6G+ cells) and monocytes/macrophages (CD11b+ Ly6G-cells) into the site of nerve injury (**Supplementary Fig. 4a and b)**. Altogether, these results indicate that although C5aR1 is expressed in resident and infiltrating leukocytes, especially monocytes/macrophages, this signaling is not involved in the recruitment of these cells into the site of nerve injury.

Infiltrated and resident leukocytes at the site of nerve injury can mediate the development of neuropathic pain through the production of a range of pro-inflammatory cytokines^12,58,59^. Thus, next we sought to verify whether C5aR1 signaling would be involved in the production of these cytokines in the site of peripheral nerve injury^31,32,34^. The increase in the production of pro-inflammatory cytokines (TNF, IL-1β, IL-6) and chemokines (CXCL1/KC CXCL2/MIP-2, CCL2/MCP-1, CCL3/MIP-1α, CCL4/MIP-1β) at the site of nerve injury was similar in *C5ar1*^-/-^ mice and WT mice (**Supplementary Fig. 4c**). These results indicate that C5aR1 signaling does not play a major role in production of pro-nociceptive cytokines at the site of peripheral nerve injury. Altogether, these data suggest that C5aR1 signaling at the site of peripheral nerve injury might be irrelevant for the development and maintenance of neuropathic pain.

### C5aR1 is expressed in sensory neurons-associated macrophages (sNAMs) of the sensory ganglia

It is well-known that neuroimmune glia interactions in the sensory ganglia also play a critical role in the development of neuropathic pain caused by peripheral nerve injury^4,10,15^. Therefore, we asked whether C5aR1 signaling on cells in the sensory ganglia would be determinant in the development and maintenance of neuropathic pain. First, we analyzed the expression of *C5ar1* in the DRGs. Conventional PCR indicated that the *C5ar1* is expressed in mice DRGs (**Fig. 3a**) and quantitative PCR showed a significant increase of *C5ar1* expression in the DRGs (L3-L5) after PSNL (**Fig. 3b)**.

**Figure 3.**
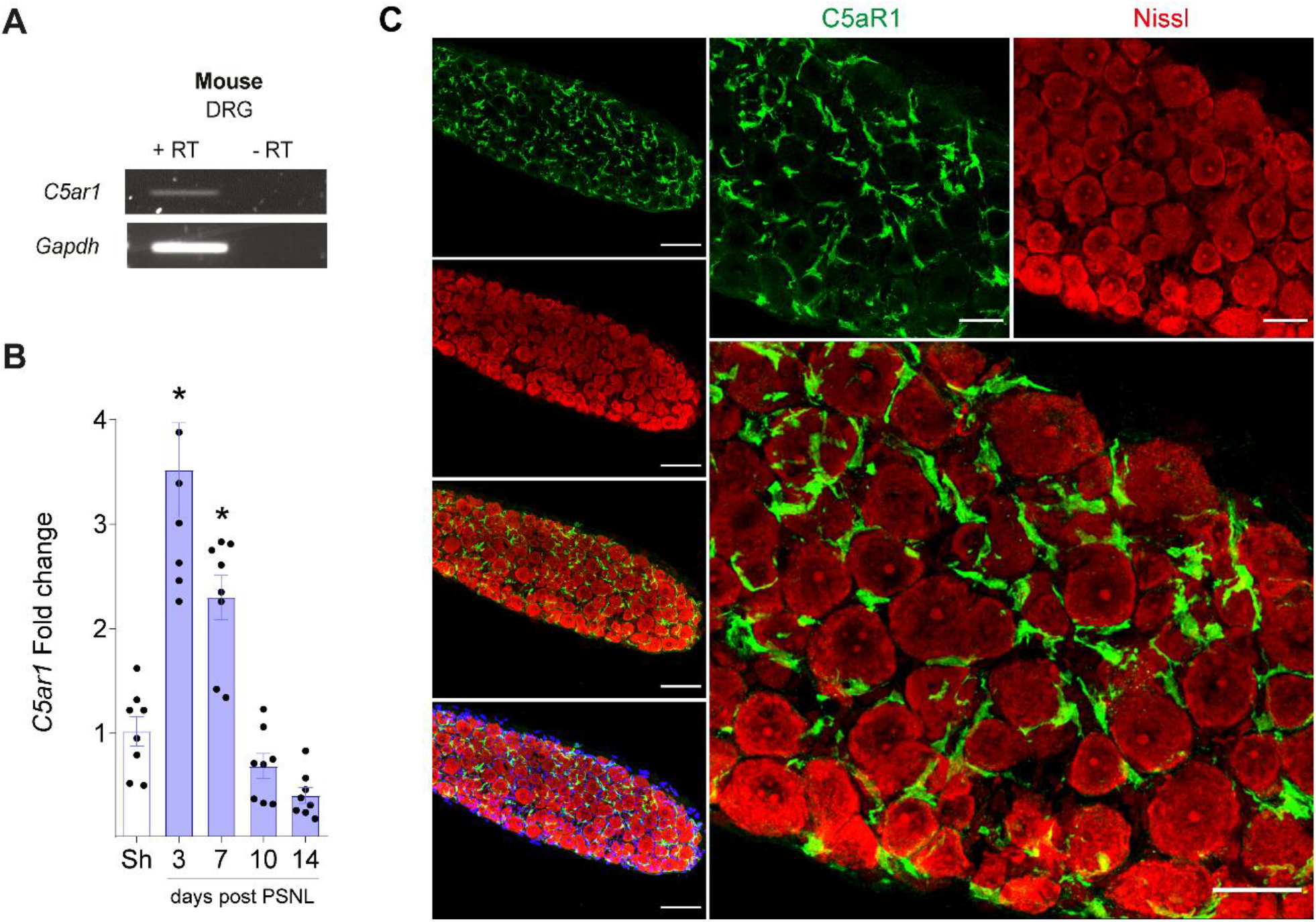
C5aR1 is expressed in the sensory ganglia and increased after PSNL. **(A)** Expression of *C5ar1* in DRG from naive mice. (**B)** Time-course of expression of *C5ar1* in the DRGs from mice after sham or PSNL. Data are represented as the mean ± s.e.m., *n*=6-8. \**P*<0.05, related to sham, analyzed by One-Way ANOVA with post hoc Bonferroni (**C)** Representative images showing immunoreactivity for C5aR1 (green color) double labeled with Nissl (red color, neuronal staining) in the DRGs of mice harvested 7 days after PSNL. Bar scale in the left panels corresponds to 100 μm. In the right panels, correspond to 40 μm.

In line with the transcriptional data, we detected a robust expression of the C5aR1 protein in the sensory ganglia after PSNL (**Fig. 3c**). Notably, C5aR1 is expressed in cells that were not labeled for Nissl staining (sensory neurons) (**Fig. 3c**). Next, we took advantage of single-cell RNA sequencing (scRNA-seq) public data set of DRGs cells to analyze the expression of *C5ar1*^52^. Notably, UMAP analysis showed that *C5ar1* is highly expressed in sNAMs of the DRGs (**Fig 4a and b**), which increased after peripheral nerve injury (Crush injury). In order to validate the scRNA-seq data, the cell profile of *C5ar1* expression in the DRGs after PSNL was further analyzed by immunofluorescence. Corroborating the gene expression profile, double immunostaining revealed that in the DRGs of PSNL mice, C5aR1 is exclusively expressed in cells expressing the macrophage markers Iba-1 (∼100%) and CX3CR1 (∼100%) (**Fig. 4c and Supplementary Fig. 5**), indicating its expression in sNAMs. Confirming the expression in sNAMs, the analysis of DRGs from *C5ar1*-GFP mice also indicated that GFP-expressing cells also express Iba-1 (**Fig. 4d**). There was no evidence of significant expression of C5aR1 in satellite glial cells (Kir 4.1 positive cells) by single cell transcriptome analysis or immunofluorescence or (**Fig 4a and b, Supplementary Fig. 6**).

**Figure 4.**
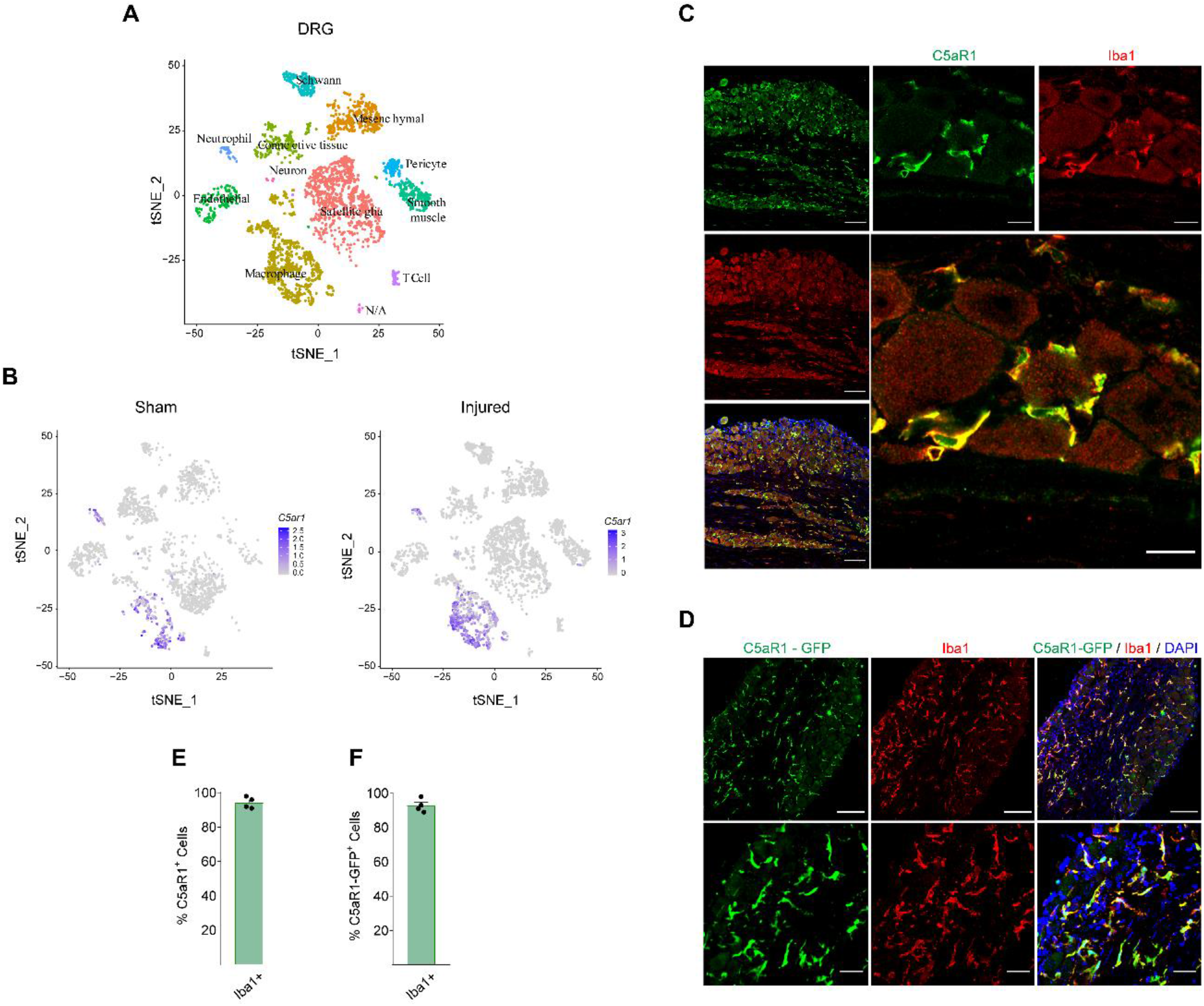
C5aR1 is expressed in the sensory neurons-associated macrophages (sNAMs) of the mouse sensory ganglia. **(A)** tSNE plot representation of Cell population encountered in the mouse DRGs by using single-cell RNAseq re-analysis (GSE139103) (**B)** tSNE plot representation showing the expression of *C5ar1* in different cell populations in the DRGs from naive and peripheral nerve injured mouse (crush injury). **(C)** Representative images and (**D)** quantification showing immunoreactivity for C5aR1 (green color) double/triple labeled with DAPI (cell nuclei; blue) and Iba-1 (red color, macrophages) in the DRGs harvested from WT mice 7 day after PSNL. Bar scale 100 μm (left panels) and 15 µm (right panels). Data are represented as the mean ± s.e.m., *n*=4. (**E)** Representative images and (**F)** quantification showing immunoreactivity for C5aR1-GFP (green color) double/triple labeled with DAPI (cell nuclei; blue) and Iba-1 (red color, macrophages) in the DRGs harvested from C5aR1-GFP mice 7 day after PSNL. Bar scale 100 μm (upper panels) and 30 µm (lower panels). Data are represented as the mean ± s.e.m., *n*=4.

To test the translational potential of our findings, we next analyzed the expression of C5AR1 in postmortem human DRG tissues from organ donors. PCR analysis showed that *C5AR1* is expressed in human DRGs (**Fig. 5a**). Furthermore, C5AR1 protein was also detected in the human DRGs, which expression was found to colocalize with the canonical maker of macrophages, Iba-1 (**Fig. 5b**). Collectively, these results indicated that C5aR1 is expressed in resident sNAMs of mouse and human DRGs.

**Figure 5.**
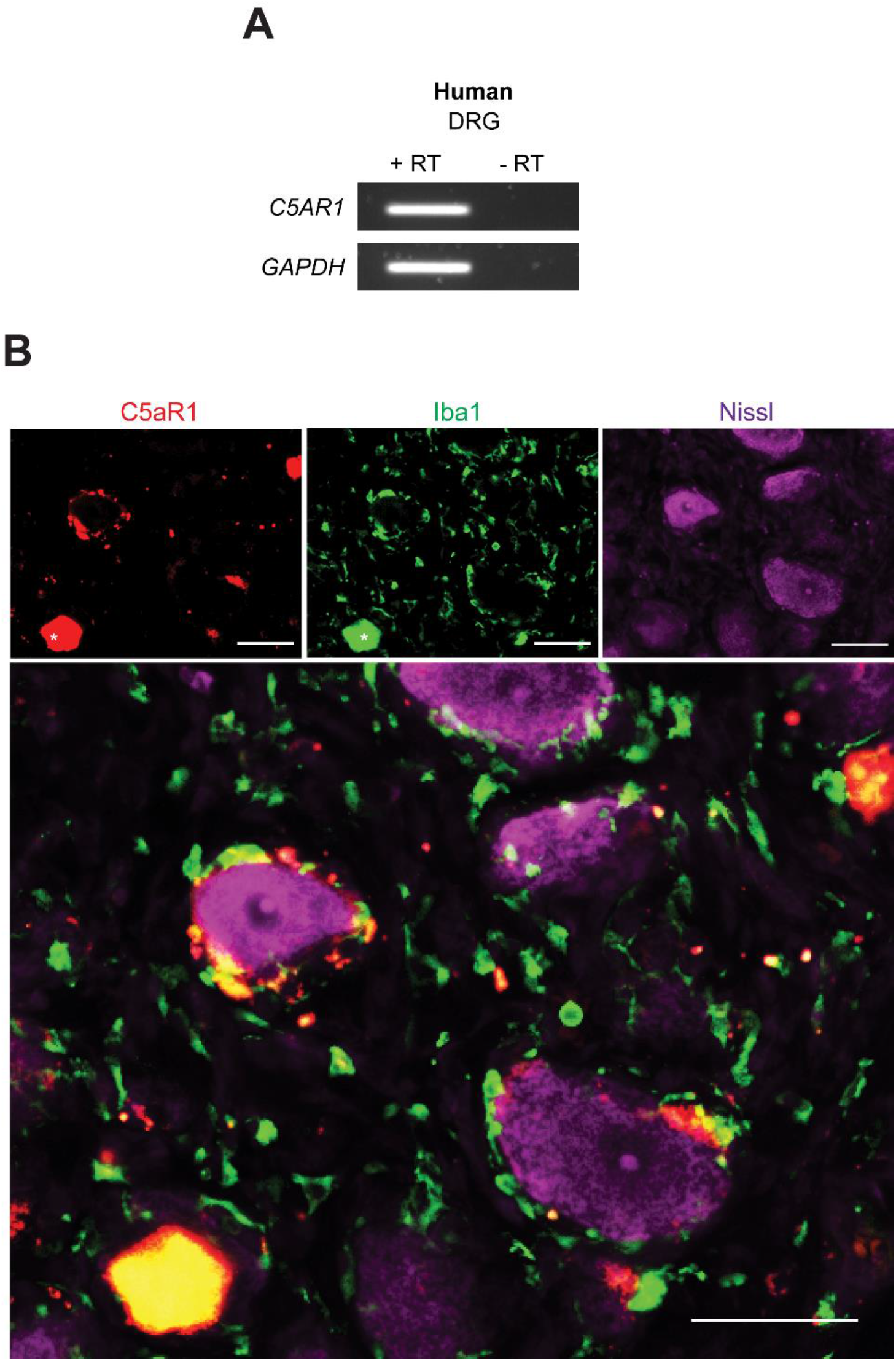
C5aR1 is expressed in sensory neurons-associated macrophages (sNAMs) of the human sensory ganglia. **(A)** *C5AR1* expression in human DRGs analysed by conventional RT-PCR **(B)** Representative images showing immunoreactivity for C5aR1 (red color) triple labeled with Iba-1 (green color, macrophages) and Nissl (purple color, neuronal staining) in the human DRGs slices Bars correspond to 50 μm.

### C5aR1 signaling in sNAMs of the sensory ganglia mediates neuropathic pain

We and others have recently identified sNAMs in the sensory ganglia (eg. DRGs) as crucial for the development of neuropathic pain caused by peripheral nerve injury^13,15,16,60,61^. Therefore, next we investigated whether C5aR1 signaling in sNAMs in the DRGs plays a role in the development of neuropathic pain. To target DRGs and not the site of injury, we delivered DF2593A intrathecally (i.th.)^62^, and assessed its effect on neuropathic pain after PSNL. We found that the i.th. treatment of mice with DF2593A reduced mechanical and cold pain hypersensitivity triggered by peripheral nerve injury (**Fig. 6a**). We next developed conditional deficient mice in which C5aR1 is not expressed in sNAMs. Two different strains of mice carrying promoters driving Cre-recombinase were used, the *Lyz2 (LysM)* promoter-dependent Cre (*Lyz2*-Cre) and *Cx3cr1* promoter- and tamoxifen-dependent Cre (*Cx3cr1*-Cre^ER^). These two strains of mice have been used for specific knockout genes in resident macrophages^39,40,63^. The *Lyz2*-Cre and *Cx3cr1*-Cre^ER^ mice were then backcrossed with *C5ar1*^flox/flox^, generating *Lyz2*-Cre^+/-^ - *C5ar1*^flox/flox^ (**Fig. 6b**), *Cx3cr1*Cre^ER+/-^ - *C5ar*1^flox/flox^ mice (**Fig. 6c**) and respective littermate controls. The use of these two lines of mice was necessary to exclude the possible contribution of C5ar1 signaling in microglia for the development of neuropathic pain. In fact, whereas the use *Cx3cr1*-Cre^ER^ allows for deletion of C5ar1 signaling on microglia and sNAMs, *Lyz2*-Cre^+/-^ has been shown to spare microglial genes^64,65^. Conditional deletion of *C5ar1* in sNAMs reduced mechanical and cold pain hypersensitivity induced by PSNL (**Fig. 6b and c**), without altering basal nociceptive behaviors (**Supplementary Fig. 7**). In order to confirm the pronociceptive role of C5aR1 signaling in sNAMs we tested the ability of recombinant C5a to promote pain hypersensitivity when injected intrathecally, which reaches the DRGs. Recombinant C5a induced mechanical and cold pain hypersensitivity in a dose-time- and C5ar1-dependent manner (**Fig. 6d**). Furthermore, mechanical and cold pain hypersensitivity triggered by recombinant C5a were also reduced in *Lyz2*-Cre^+/-^ - *C5ar*1^flox/flox^ mice compared to littermate controls (**Fig. 6e**). These results together indicate that sNAMs in the sensory ganglia play a critical role in the development of neuropathic pain caused by peripheral nerve injury.

**Figure 6.**
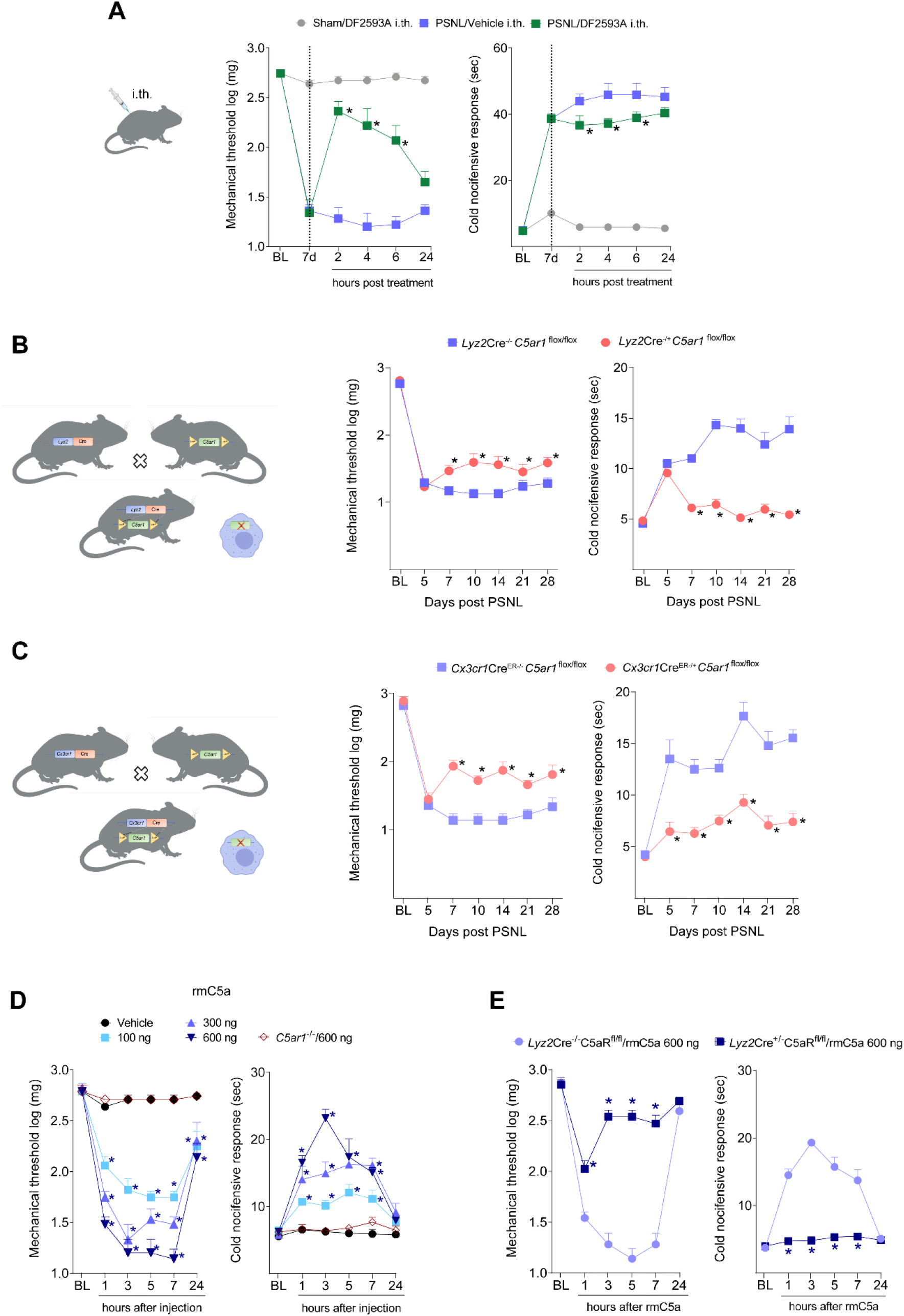
C5aR1 signaling in sNAMs of the sensory ganglia drives neuropathic pain. (A) Intrathecal treatment of WT PSNL mice with C5aR1 antagonist (DF2593A, 30 μg/5 μl) or vehicle 7 days after PSNL. Mechanical and cold allodynia were determined before and up to 24 h after treatment. PSNL was induced in (B) Lyz2-Cre-C5ar1flox/flox, (C) Cx3cr1CreERC5ar1flox/flox mice or in control littermates. Mechanical and cold allodynia were determined before and up to 28 days after PSNL. (D) WT and C5ar1-/-mice received an intrathecal injection of rmC5a (100-600 ng/5 μl). Mechanical and cold allodynia were determined before and up to 24 h after rmC5a injection. (E) Lyz2-Cre-C5ar1flox/flox and control littermates (600 ng/5 μl) mice received an intrathecal injection of rmC5a (600 ng/5 μl). Mechanical and cold allodynia were determined before and up to 24 h after rmC5a injection. Data are represented as the mean ± s.e.m., n=6-8. *P<0.05 compared to the respective control groups and analyzed by repeated measures Two-Way ANOVA with Bonferroni post hoc.

### C5aR1 signaling in sNAMs mediates neuropathic pain through IL-1β

Activated sNAMs mediate the development of neuropathic pain through the production of pro-inflammatory cytokines, especially TNF and IL-1β^15,16^. Thus, we investigated whether C5aR1 signaling in sNAMs mediates neuropathic pain through the production of pro-inflammatory cytokines. First, we analyzed the expression of makers of sNAMs and SGCs activation (*Aif1* and *Gfap, respectively*) and cytokines (*Tnf* and *Il1b*) in the DRGs from WT and *C5ar1*^-/-^ mice after PSNL. We found that whereas the expression of *Gfap, Aif1* and *Tnf* was not changed, *Il1b* expression was reduced in the DRGs from *C5ar1*^-/-^ mice compared to WT mice after PSNL (**Fig. 7a**). The *Il1b* expression in the DRGs after peripheral nerve injury was also reduced in *Lyz2*-Cre^+/-^-*C5ar*1^flox/flox^ mice compared to littermate controls (**Fig. 7b**). Supporting these findings, the intrathecal injection of recombinant C5a also increased the expression of *Il1b* in the DRGs from WT mice, which was abrogated in the DRGs from *C5ar1*^-/-^ mice (**Fig. 7c**). Furthermore, the ability of recombinant C5a to trigger the expression of *Il1b* was almost completely abrogated in *Lyz2*-Cre^+/-^ - *C5ar*1^flox/flox^ mice compared to littermate controls (**Fig. 7d)**. Finally, confirming the hypothesis that C5aR1 signaling in sNAMs mediates neuropathic pain through IL-1β, the pronociceptive activity promoted by intrathecal injection of recombinant C5a was reduced in *Il1r1* null (*Il1r*^-/-^) mice (**Fig. 7e**). Together, these results indicate that C5aR1 signaling in sNAMs of the sensory ganglia mediates neuropathic pain via stimulation of IL-1β production (**Supplementary Fig. 8**).

**Figure 7.**
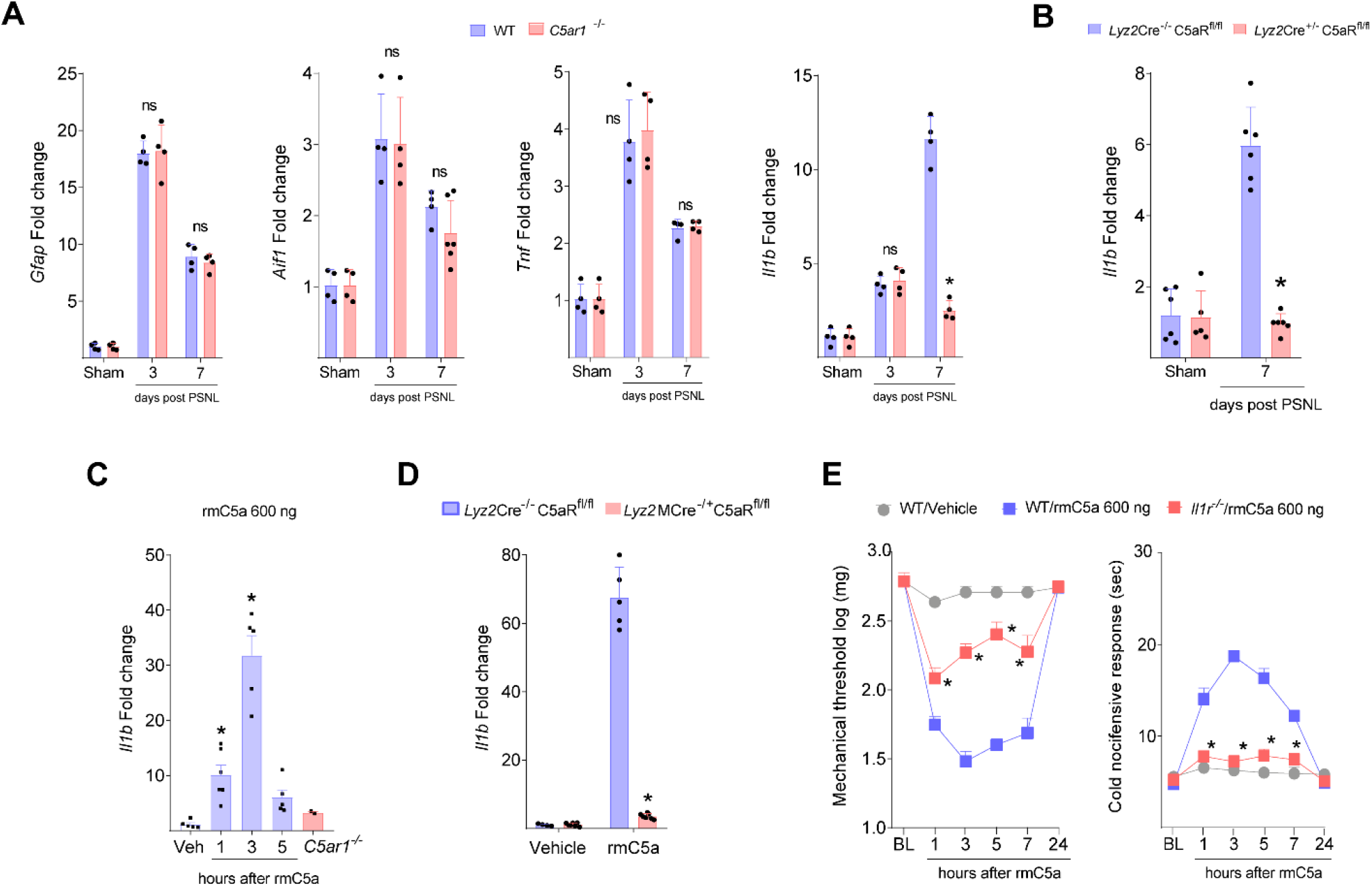
C5aR1 signaling in sNAMs mediates neuropathic pain through IL-1b. **(A)** mRNA expression of *Gfap, Aif1, Tnf and IL-1b* in the ipsilateral DRGs from WT or *C5ar1*^-/-^ mice at indicated time points after PSNL or sham surgery, *n*=4. **(B)** mRNA expression of *IL-1b* in the ipsilateral DRGs from *Lyz2*-Cre-*C5ar*1^flox/flox^ mice or control littermates 7 days after PSNL or sham surgery, *n*=5-6. (**C)** Time-course (hours) of mRNA expression of *IL-1b* in the DRGs from WT or *C5ar1*^-/-^ mice after intrathecal injection of rmC5a (600 ng/5 μl) or vehicle, *n*=5-6. mRNA expression of *IL-1b* in the DRGs from *Lyz2*-Cre-*C5ar*1^flox/flox^ mice or control littermates 3 h after intrathecal injection of rmC5a (600 ng/5 μl), *n*=5-6. **(E)** WT and *Il1r1*^*-/-*^ mice received an intrathecal injection of rmC5a (600 ng/5 μl). Mechanical and cold allodynia were determined before and up to 24 h after rmC5a injection, *n*=6. Data are represented as the mean ± s.e.m. analyzed by One-Way ANOVA (A-D) or Two-Way ANOVA (E), with Bonferroni post hoc test. \**P*<0.05 compared to vehicle or littermate controls.

## Discussion

Although pain is a distressing sensation, often it is temporary and beneficial^1^. However, chronic condition, which is associated with conditions like neuropathy is particularly refractory to the existing treatments, which mostly target neuronal pathways. Growing evidence points towards neuro-immune glial interactions, including C5a/C5aR1 signaling^19,36,37^, as main players in the development and maintenance of neuropathic pain, and therefore a potential novel therapeutic target^4–10^. Herein we focus upon the loci and mechanisms by which C5a/C5aR1 signaling drives peripheral nerve injury-induced neuropathic pain. We demonstrate for the first time the role of sensory neurons-associated macrophages (sNAMs) in the sensory ganglia, but not in macrophages of the site of peripheral nerve injury and that this signaling triggers the production of IL-1β, thereby mediating the induction of pain hypersensitivity.

Consistent with previous studies we confirmed that C5a/C5aR1 signaling plays an important role in neuropathic pain responses following peripheral nerve injury^36,37^ in male and female mice, despite gender differences in neuro-immune mechanisms^53^. The phenotype of *C5ar1* null male and female mice in PSNL-induced neuropathic pain suggests that blockade of C5a/C5aR1 signaling is a new target for the prevention of neuropathic pain development. We confirmed this through the demonstration that the novel potent and selective orally-active allosteric antagonist of C5aR1 (DF2593A)^37^ given during the development of neuropathic pain, or even when the neuropathic pain is established, significantly reduced mechanical and cold pain hypersensitivity induced by PSNL.

The contribution of neuroimmune glia interactions in sensory ganglia during the development of neuropathic pain are established^4,9,39^. The production of pronociceptive cytokines, especially TNF and IL-1β, by satellite glial cells and macrophages in the DRGs is a well-reported mechanism underlying the development of neuropathic pain^15,16,61,67^. However, the signaling mechanisms driving the release of these molecules had yet to be identified. Herein we demonstrate that C5a/C5aR1 signaling in sNAMs triggers the production of the pro-nociceptive IL-1β but not TNF. IL-1β has multiple mechanisms of action which synergize to increase neuronal excitability and therefore identification of the key process driving the generation of this molecule could be a critical breakpoint in this process. IL-1β acts directly to sensitize sensory neurons^66^ via IL-1R1 which are expressed by them^66^. *In vitro*, IL-1β enhances DRG neuronal excitability by altering the expression of receptors such as TRPV1, sodium channels, GABA receptors, and NMDA receptors^45,66^. Conditional deletion of IL-1R1 in sensory nociceptive neurons reduces pathological pain^66^ and IL-1β stimulates the production of pro-inflammatory and pro-nociceptive mediators such as Brain Derived Neurotrophic Factor^16^, which also contribute to the development of neuropathic pain.

Although we identified an abundance of C5aR1-expressing monocytes/macrophages at the site of PSNL injury, neither the recruitment of immune cells (macrophages and neutrophils) to the site of PSNL injury nor the production/release of pro-nociceptive cytokines and chemokines were altered in *C5ar1* null mice. It is likely that due to a redundancy in signaling arising from the multiple chemotactic substances released at the site of nerve injury, which can attract these leukocytes and stimulate cytokines/chemokines production, C5aR1 mechanisms are not critical at this site. This finding is corroborated by the recent report which ruled out participation of macrophages/monocytes at the site of peripheral nerve injury in neuropathic pain development^16^. We have confirmed that in the sensory ganglia, C5aR1 is exclusively expressed in sNAMs, and that this expression increased following peripheral nerve injury. Importantly, the translational relevance of these murine data we confirmed by the demonstration that C5AR1 is present in human sNAMs. Using multiple complementary approaches, we provide evidence that C5aR1 signaling in sNAMs contributes to the development of neuropathic pain. Firstly, intrathecal delivery of C5aR1 antagonist that target sNAMs, but not peripheral cells, reduced established neuropathic pain. The involvement of C5aR1 in sNAMs in the development of neuropathic pain was corroborated by using two different strains of C5aR1 conditional mice. It is important to note that there are no sNAM specific genes that could be used as a driver of Cre-recombinase to knockout *C5ar1* gene only in these cells. Therefore, we selected a combination of *Lyz2* and *Cx3cr1 genes promoters*, both of which are expressed by sNAMs, but not by the local infiltrating monocytes (Lyz2+ CX3CR1-)- nor microglia (Cx3cr1+ Lyz2-). The reduction of pain hypersensitivity in both conditional strains compared to littermate controls further support the hypothesis that C5aR1 signaling on sNAMs play a role in neuropathic pain development. Nevertheless, we cannot exclude that C5aR1 signaling on microglia might be involved in this process as already suggested^36^

In summary, our results reveal the mechanism by which C5a/C5aR1 signaling mediates the development of neuropathic pain. C5a/C5aR1 signaling in sNAMs of the sensory ganglia, but not at the site of nerve injury or spinal cord, and drives neuropathic pain through the stimulation of IL-1β expression. These results with the observation of the C5aR1 in human sNAMs provide strong evidences that C5a/C5aR1 signaling is a promising translational target for the clinical treatment of neuropathic pain.

## Materials and Methods

### Animals

The experiments were performed in *C5ar1* deficient mice (*C5ar1*^-/-^, 20-25g, Jackson Laboratories, Stock number 6845, Strain *C5ar1*^*tm1Cge*^)^38^ and respective controls (Balb/C wild type, WT, 20-25 g). Also, C57Bl/6 mice (wild type, WT, 20-25 g), paired with *C5ar1*^flox/flox^-GFP and *Lyz2-Cre*^+/-^-C5aR^Flox/Flox^ mice were generated as described previously^39^. We also used a heterozygous mice that express *Cx3cr1*^GFP/+^ (20–25g)^40^ and a *Cx3cr1*-Cre^ER^ (20–25g, Jakson Laboratories, Stock number 021160, Strain *Cx3cr1*^tm2.1(cre/ERT2)41^.

For all experiments involving removal of sNAMs *C5ar1*, the *CX3CR1*^*CreER/+*^ line mice were crossed to *C5ar1*^flox/flox^ mice. Conditional and littermate controls, 6-8 weeks old, were treated twice with 4 mg tamoxifen (Sigma) diluted in 200 µL of corn oil (Sigma), injected subcutaneously at two time points 48 h apart.

WT (Balb/C and C57Bl/6) strains were obtained from the Animal Care Facility of Ribeirão Preto Medical School, University of São Paulo. Local colonies of transgenic mice were established and maintained at the Animal Care Facility of Ribeirão Preto Medical School, University of São Paulo. Food and water were available *ad libitum* and controlled light-dark cycle. Animal experiments were performed in accordance with the International Association for the Study of Pain guidelines^42^ for those animals used in pain research and they were approved by the Committee for Ethics in Animal Research of the Ribeirao Preto Medical School -USP (Process n° 120/2014).

### Drugs

DF2593A^37^ was gently given by Dompé Biopharmaceutical and diluted in saline at the moment of use. Mice received C5a recombinant (R&D Systems; Cat. 2150), diluted in BSA 0.1% and stocked at -70 °C until the moment of use.

### Neuropathic pain model (PSNL – partial sciatic nerve injury)

Partial sciatic nerve ligation (PSNL) was performed as previously decribed^11,43,44^. Under anesthesia with 2% isoflurane, the animal muscle was cut in sterile conditions. The nerve was isolated with a glass pipette, and a silk suture 7.0 (Ethicon, Johnson & Johnson, Cat. 7733, needle 3/8 6.5 mm) was inserted into the nerve, which was tightly ligated, so that 1/3 to 1/2 of the nerve thickness was trapped in the ligature. The wound was closed with a 4.0 silk suture (Ethicon, Johnson & Johnson, Cat. D2764, needle 1/2 17 mm).

### Behavioral nociceptive tests

All behavioral tests were performed after a minimum 30 minutes of mice habituation to the environment and assays.

### von Frey filament test

For testing mechanical nociceptive threshold, mice were placed on an elevated wire grid and the plantar surface of the ipsilateral hind paw stimulated perpendicularly with a series of von Frey filaments (North Coast Medical), with logarithmically increasing stiffness. Each one of these filaments was applied for approximately 5 seconds to induce a paw-withdrawal reflex. The weakest filament able to elicit a response was taken to be the mechanical withdrawal threshold. The log stiffness of the hairs is determined by log10 (milligrams)^11,45,46^.

### Electronic von Frey test

In the same elevated wire grid described, a hind paw flexion reflex was evoked with a hand-held force transducer, adapted with a 0.5-mm2 polypropylene tip (Insight, Brazil). The stimulus was applied in the hind paw, with a gradual increase in pressure. When the paw was withdrawn, the stimulus was automatically discontinued and its intensity recorded. The end point was characterized by the removal of the paw in a clear flinch response after the paw withdrawal^47^.

### Acetone test

Mice were habituated in the same cages and elevated grids used during the von Frey tests. Then, a drop (50 µL) of acetone was applied to the plantar surface of the hind paw using a syringe of 1 ml. The time spent with nociceptive behaviors - defined as flinching, licking or biting the limb - were timed within 60 seconds after the application of acetone^11,46,48^.

### Hargreaves test

The latency of paw withdrawal to radiant heat stimuli was measured using the Hargreaves apparatus (Ugo Basile, Italy) and described previously^37,46^. Mice were habituated in an elevated glass surface of constant temperature (37°C). An infrared light source was applied perpendicular on the plantar surface of each mouse’s hind paw. The end point was characterized by the removal of the paw followed by clear flinching movements. Latency (time in seconds) to paw withdrawal was automatically recorded. To avoid tissue damage, a maximum latency (cut-off) was set at 20 seconds.

### Hot-Plate Test

The noxious heat thresholds were also examined using the Hot-Plate test (Ugo Basile, Italy). Mice were placed in a 10-cm-wide glass cylinder on a hot plate maintained at 48°C, 52°C or 56°C. Two control latencies at least 20 minutes apart were determined for each mouse. The latencies in seconds for hind paws licking or jumping for each animal were recorded. To avoid tissue damage, a maximum latency (cut-off) was set at 20 seconds^37,46^.

### Rotarod test

The apparatus consisted of a bar with a diameter of 2.5 cm, sub-divided into five compartments by disks 25 cm in diameter (Ugo Basile, Italy). The bar rotated at a constant speed of 22 rotations per minute. The animals were selected 24 hours previously by eliminating those mice that did not remain on the bar for two consecutive periods of 120 seconds. Mice were submitted to walk on the rotate cylinder for a total of 120 seconds. The time that each animal remained walking was registered as latency to fall^37,46^.

### Drugs Administration

#### Oral administration

Animals were manually restricted, without anesthesia. The drugs were injected orally, 10 µl/10 g of body weight, with a metal gavage curve needle and a 1 mL syringe. Acute treatments were made after 12 hours of food restriction and free water consumption.

#### Intrathecal injection

Under isoflurane (2%) anesthesia, mice were securely hold in one hand by the pelvic girdle and inserting a BD Ultra-Fine® (29G) insulin syringe directly on subarachnoid space (close to L4–L5 segments) of the spinal cord. A sudden lateral movement of the tail indicated proper placement of the needle in the intrathecal space^49^. For all deliveries, we used 5 µL of volume. After delivery, the syringe was held in the specific position for a few seconds and progressively removed to avoid any outflow of the substances. In all experiments in which intrathecal injections (i.t.) were performed, the vehicle was replaced by aCSF (artificial cerebrospinal fluid) (Tocris, Cat. 3525).

#### Mice tissue samples collection

Under injected anesthesia (50% xylazine, 50% ketamine – 10 µL/g body weight), animals were transcardiacally perfused with PBS buffer at room temperature (20 mL/min for 2 minutes) to wash the blood and consequently the circulating leukocytes. For immunofluorescence assays, an additional perfusion time of 5 minutes with ice PFA (Sigma Aldrich, Cat. 158127) 4% was performed (also 20 mL/minute). An incision was made in the lumbar region and the skin and muscle were removed. After a laminectomy, the DRG or spinal cord segment corresponding to L3-L5 were removed. For nerve collection, the ipsilateral thigh was cut and a fragment of approximately 1 cm of sciatic nerve was removed (0.5 cm medial and 0.5 cm lateral to ligation).

#### Human samples

Analyses were also performed on non-diseased human lumbar dorsal root ganglia that were recovered from consented and deidentified organ donors in collaboration with the LifeCenter at Cincinnati or obtained from donors through National Disease Research Interchange (NDRI) with permission of exemption from Institutional Review Board (IRB) of the University of Cincinnati.

#### ELISA (Enzyme-Linked Immunosorbent Assay)

Sciatic nerve samples, once collected, were immediately stored on ice dry and frozen at -70 °C. For processing, they were diluted in a PBS buffer, containing a protease inhibitor cocktail (Complete® Roche, Cat. 11697498001) and 0.2% of Triton X^50^. The amount of total protein in each sample was measured by the BCA method. Concentrations of cytokines were determined according to the protocol previously described^51^, using DuoSet enzyme immunoassay kits from R&D Systems.

Briefly, after grinding, the samples were centrifuged and the supernatant was incubated overnight in 96 wells plates, previously sensitized with the primary antibody of interest and with nonspecific proteins blocked by incubation with 1% BSA for 1 hour 37°C. Posteriorly, the plate was incubated at room temperature for 2 hours, with biotinylated polyclonal antibodies and then labeled with avidin-HRP for 1 hour, followed by protein-antibody-avidin/HRP link revealed by colorimetric reaction with TMB (3,3 ‘, 5,5’-tetramethylbenzidine - SureBlue - KPL, Cat. 95059). The reaction was interrupted with H2SO4 1M and the absorbance was determined at 450 nm in a SpectraMax spectrophotometer (Molecular Devices, USA).

#### Conventional RT-PCR

cDNA was synthesized with High Capacity RNA-to-cDNA Kit (Thermo Fisher Scientific, Cat. 4368814) from mouse and human dorsal root ganglia. cDNA was diluted 2:100 and used as a template for PCR experiments. PCR was performed in 20 mL of PCR buffer containing 300 nmol/L primers, 6 mL of diluted template, and 10 mL 2X Hot Start Master Mix (Apex Bioresearch Products; 42-143) for 35 cycles. The protocol included an initial 5 minutes denaturing step at 94°C, followed by 35 cycles of 30 seconds of denaturation at 94°C, 30 seconds of annealing at 56°C, and 1min of elongation at 72°C. The reaction was completed with 10 min of final elongation. The primer pairs used are described in Supplementary table 1.

#### Real-time RT-PCR

The samples, once collected, in a Trizol reagent (Invitrogen, Cat. 15596018), were immediately stored on dry ice and later in a freezer at -70°C. After grinding the tissue, the extraction of the total RNA was carried out according to the reagent manufacturer, namely: separation in chloroform, precipitation in isopropanol (overnight -20°C) and washing with 80% ethanol. The RNA concentration of each sample was determined by the optical density at the wavelength of 260 nm (NanoDrop, ThermoFisher, USA). One μg or 500 ng of total RNA was transcribed to complementary DNA by the action of the MultiScribe® reverse transcriptase enzyme, high capacity kit (Life-Invitrogen, Cat. 4368814). The amplification of the genes of interest occurred from the use of a pre-delineated IDT® primer for *C5ar1* and the endogenous GAPDH control. For the other targets, Sigma standard primers were used, the sequence of which is described in Supplementary table 2. RT-PCR was performed with a final reaction volume of 10 µL, SYBR-Green. The reaction was performed using the StepOne Real-Time PCR System (Life-Applied Biosystems). Results were analyzed using the comparative method of “cycle threshold” (CT).

#### Immunofluorescence

The samples, once collected, were immediately stored in cold 4% PFA and kept at 4°C for 4 hours and then transferred to the 30% sucrose solution for 2 days at 4°C or until sink. On dry ice, tissues were embedded in TissueTek® / OCT (Sakura) blocks and cut on cryostat: 14 μm for DRG and sciatic nerve, collected on gelatinized slides and 40 μm for spinal cord, collected in 50 ml tube containing PBS for free floating immunofluorescence. In plate or slide, the samples were washed with a PBS buffer, incubated in blocking solution (1% BSA, 22.52 mg/mL glycine in PBST - PBS+ 0.1% Tween 20), both at room temperature. Posteriorly, tissues were incubated overnight with the primary antibody at 4°C and after, for 2 hours with the secondary antibody coupled to fluorophore. The list of antibodies used is described in Supplementary table 3. The slides were assembled using VectaShield® with DAPI (Vector, Cat. H-1200). Images were acquired using a confocal Zeiss fluorescence microscope (AxioObserver LSM780) or a Keyence microscope (BZ-X800) and images were analyzed using NIH Image J open source software.

#### Flow cytometry acquisition and analysis

Sciatic nerve was collected from naïve, or PSNL mice and incubated in a solution containing 1 ml of DMEM medium and 2 mg/ml of collagenase type II (Sigma-Aldrich) for a 1 h at 37°C. After digestion, Dorsal Root Ganglia and Sciatic nerve was mechanically grinded through 40 μm cell strainers and the cell suspension was washed with PBS. The cells obtained were then washed and stained for 15 min at room temperature with fixable viability dye (L34976, Invitrogen, 1:5000) for dead cells exclusion and fluorochrome-labeled monoclonal antibodies against surface cell markers. It was used the following monoclonal antibodies: anti-CD45-BV421 (30-F11, BD Biosciences, 1:350), anti-LYG6 -APC (RB6-8C5, BD Biosciences, 1:250), anti-CD11b-PE (M1/10, BD Biosciences, 1:250). Flow cytometry analyses were performed using a FACSVerse instrument (BD Biosciences, San Jose, CA, USA) and data analyzed by FlowJo software (Treestar, Ashland, OR, USA).

#### Re-analysis of public scRNA-seq data

The single cell RNA-sequencing data from mice DRGs was acquired from the Gene Expression Omnibus (GEO) database under the series number GSE139103^52^. The single cell libraries were generated using GemCode Single-Cell 3′ Gel Bead and Library Kit on the 10X Chromium system (10X Genomics). The dataset contains cells from four animals, of which two naive mice and two with injured DRG. The feature barcode matrix was analyzed using Seurat v3. The cells were filtered according to the criteria: 600-10000 total reads per cell, 500-4000 expressed genes per cell and mitochondrial reads <10%. Clusters were identified using shared nearest neighbor (SNN) based clustering based on the first 30 PCAs and resolution = 0.5. The same principal components were used to generate the t-SNE projections. Differentially expressed genes between samples from naive and injured mice for each cluster were identified using FDR < 0.05 and |avg_log2FC| > 0.25.

#### Data analyses and statistics

Data are reported as the means ± S.E.M. The normal distribution of data was analyzed by D’Agostino and Pearson test. Each statistical test is indicated in the figure legend of each experiment. *P* values less than 0.05 were considered significant. Statistical analysis was performed with GraphPad Prism 8 (GraphPad Software, USA).

#### Data availability

The data supporting the findings of this study are available within the article and its Supplementary Information files or from the corresponding authors upon request.

## Acknowledgments

The authors gratefully acknowledge the technical assistance of Ieda R. Schivo, Ana Katia dos Santos, Jenna M. Turner, Sergio R. Rosa, Diva A. Sousa, Eleni Tamburus, Marcella Daruge Grando and Juliana Abumansur.

## Author Contributions

A.U.Q. designed, performed experimental work, analyzed data and prepared the manuscript. C.E.A.S. and M.D.F. performed experimental work related to FACS and analyzed data. A.G.M.M., A.H.P.L. and M.C.M.C. performed experiments related to surgery and harvest tissue. S.H.L. performed experiments with human samples. S.D. performed the single-cell analysis. JK developed and provided the *C5ar1*^*Flox/flox*^ mice and discussed the manuscript. D.R.S. performed experiments and important scientific comments. L.B., M.A., T.B., V.C., J.C.A.F., and F.Q.C., provided critical materials and comments. T.M.C designed, directed and supervised the study, interpreted data and wrote the manuscript. All authors reviewed the manuscript and provided final approval for submission.

