## Supplemental for "Complement receptor C5aR1 signaling in sensory neuron-associated macrophages drives neuropathic pain"

**This file includes:**

Figures S1 to S8
Tables S1 to S3

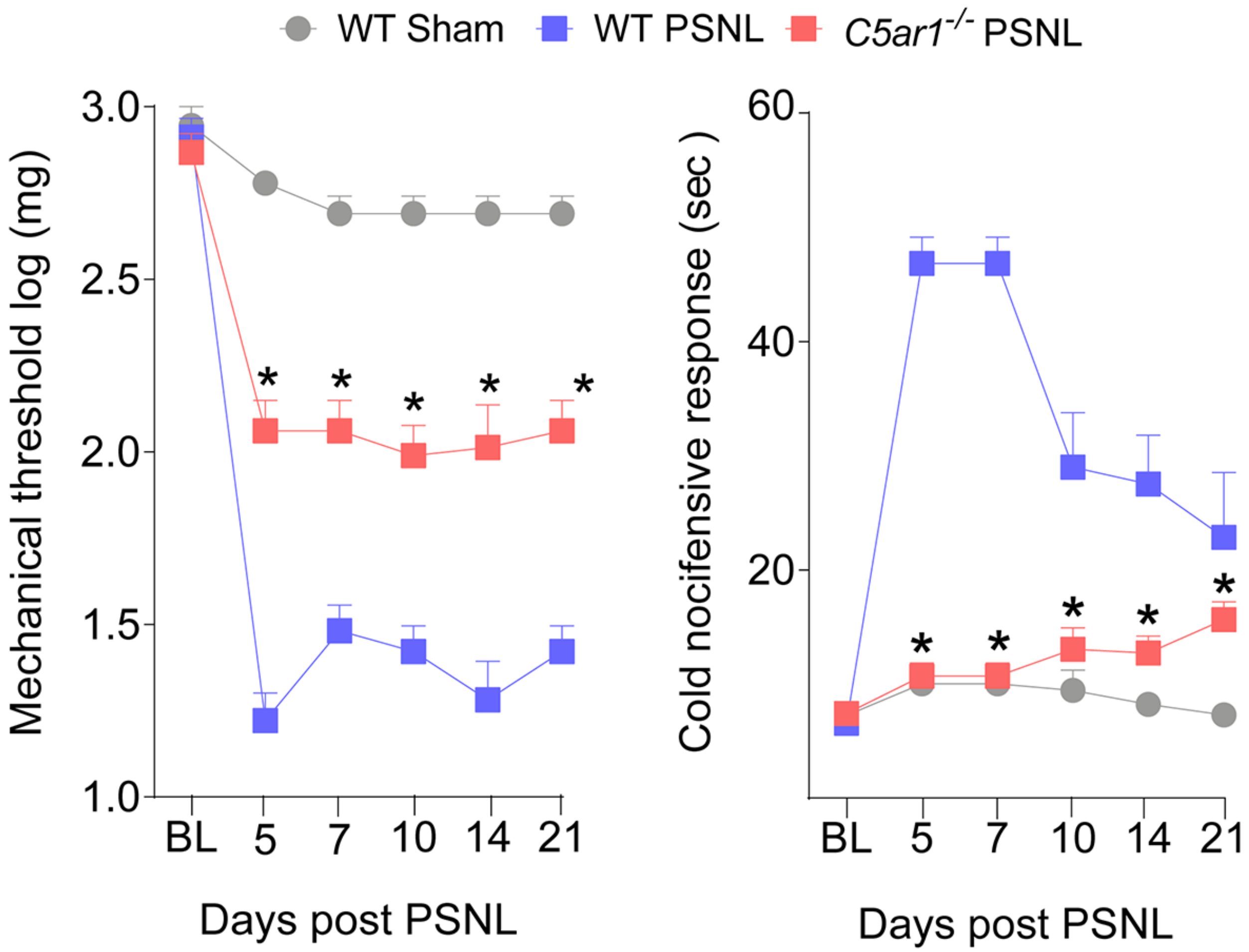


**Figure S1. C5aR1 is crucial for neuropathic pain development in female mice.** Mechanical and cold pain hypersensitivity in Balb/C WT or *C5ar1*$^{-/-}$ female mice, from 3 up to 21 days after PSNL surgery. Data represented mean ± s.e.m. and analyzed by repeated measures Two-Way ANOVA with post hoc Bonferroni, *n*=6. *$P$<0.05 compared to WT/PSNL group.

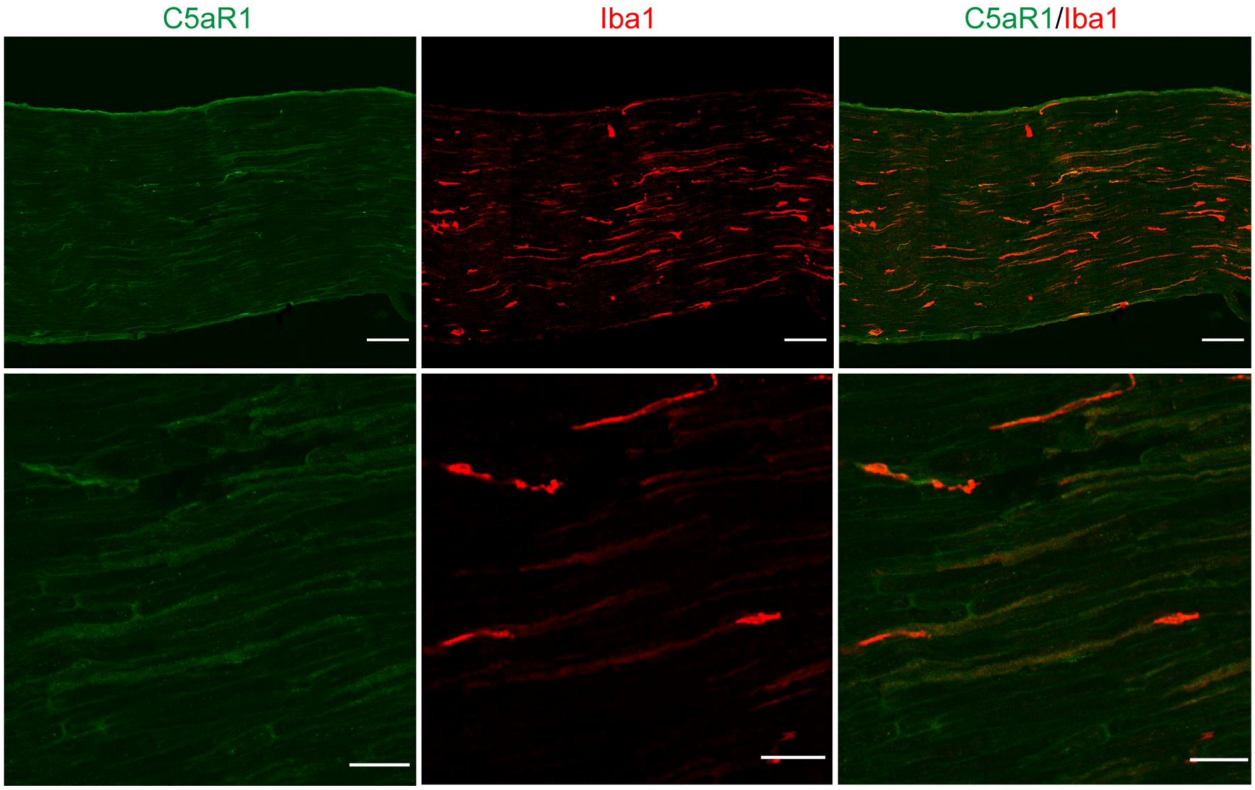


**Figure S2. Specificity of anti-C5aR1 antibody**. Representative immunostaining for C5aR1 (green) in sciatic nerve from naïve *C5ar1*$^{-/-}$ mice. Iba-1 (red) staining was used as a positive control of the IF assay and tissue integrity. Bars in upper panels correspond to 100 µm and lower panels to 30 µm.

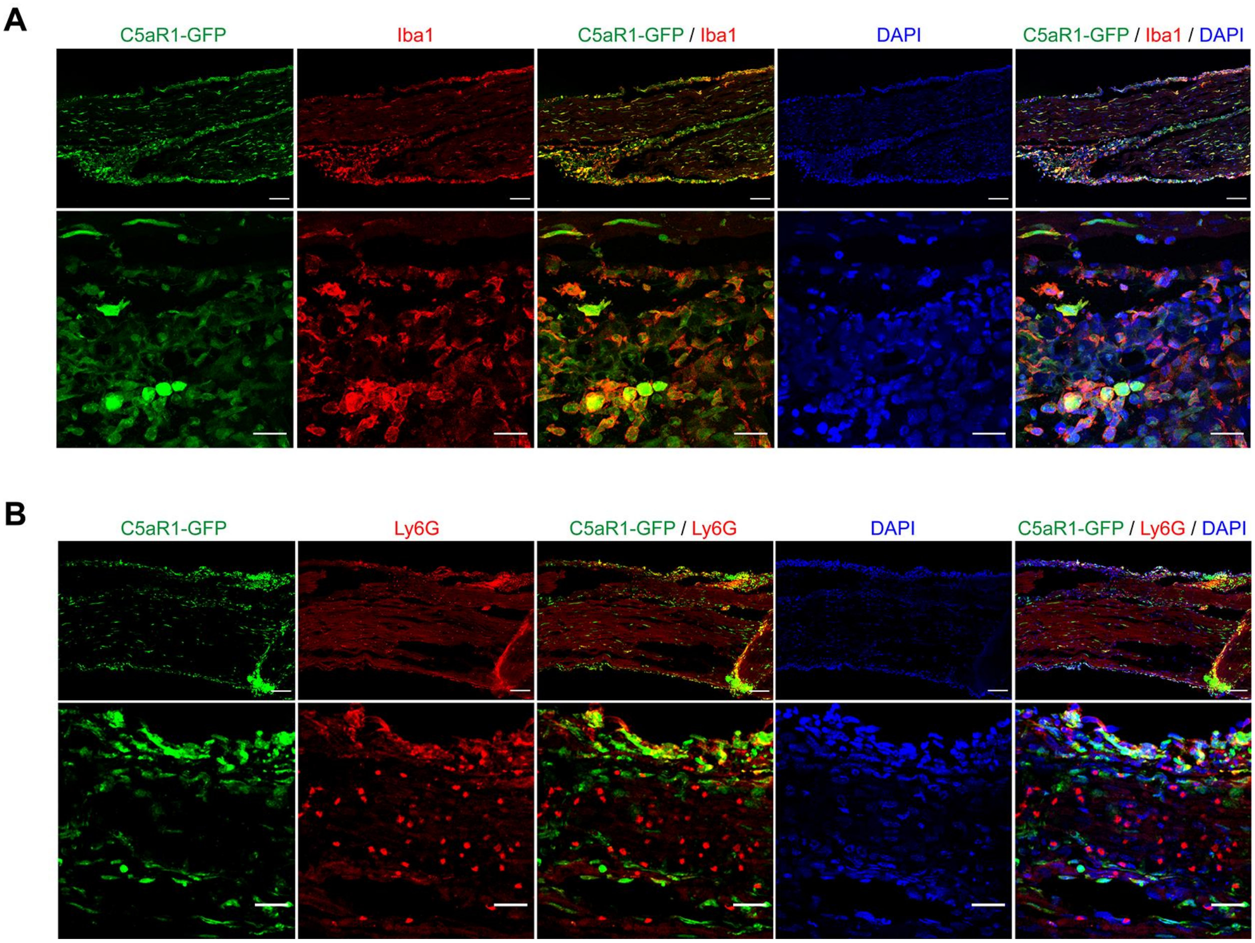


**Figure S3. C5aR1-GFP is co-expressed with macrophages marker and is detected in few Ly6G+ cells (neutrophils) in PSNL injury**. Immunofluorescence for **A.** Iba1 (red, macrophages) or **B.** Ly6G+ cells (red, neutrophils) in the sciatic nerve samples from C5aR1-GFP (green) mice at 3 days after PSNL. Bars in upper panels correspond to 100 µm and lower panels to 30 µm.

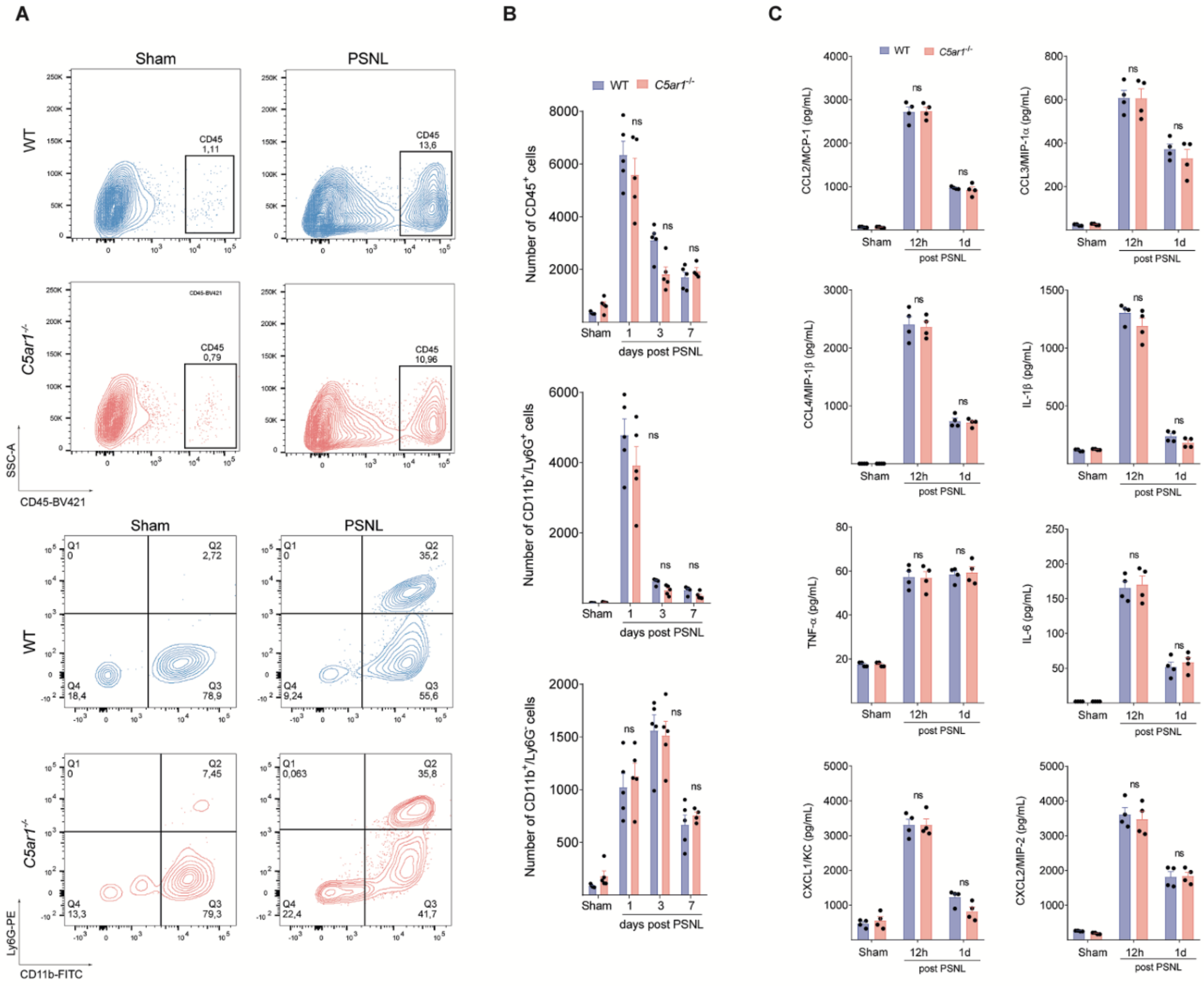


**Figure S4. C5aR1 signaling is not required for leukocyte migration or cytokine/chemokine production at the site of nerve injury.** A. Representative dot plots and B. quantification of absolute number of CD45+ cells (Leukocytes, upper panels), CD11b+ Ly6G+ cells (neutrophils) or CD11b+/Ly6G- cells (macrophages/monocytes, bottom panels) in the sciatic nerve of WT or C5ar1-/- mice harvested from naive or PSNL (n=5). Data are represented as the mean ± s.e.m. and analyzed by repeated measures One-Way ANOVA with post-hoc Bonferroni. *P<0.05 compared to sham; ns-non-significant C. WT or C5ar1-/- mice sciatic nerve was harvested from naive or PSNL mice at indicated time points. The levels of cytokines/chemokines were measured by ELISA. Data are represented as the mean ± s.e.m. and analyzed by One-Way ANOVA, with Bonferroni post hoc test , n=4. ns, non significant.

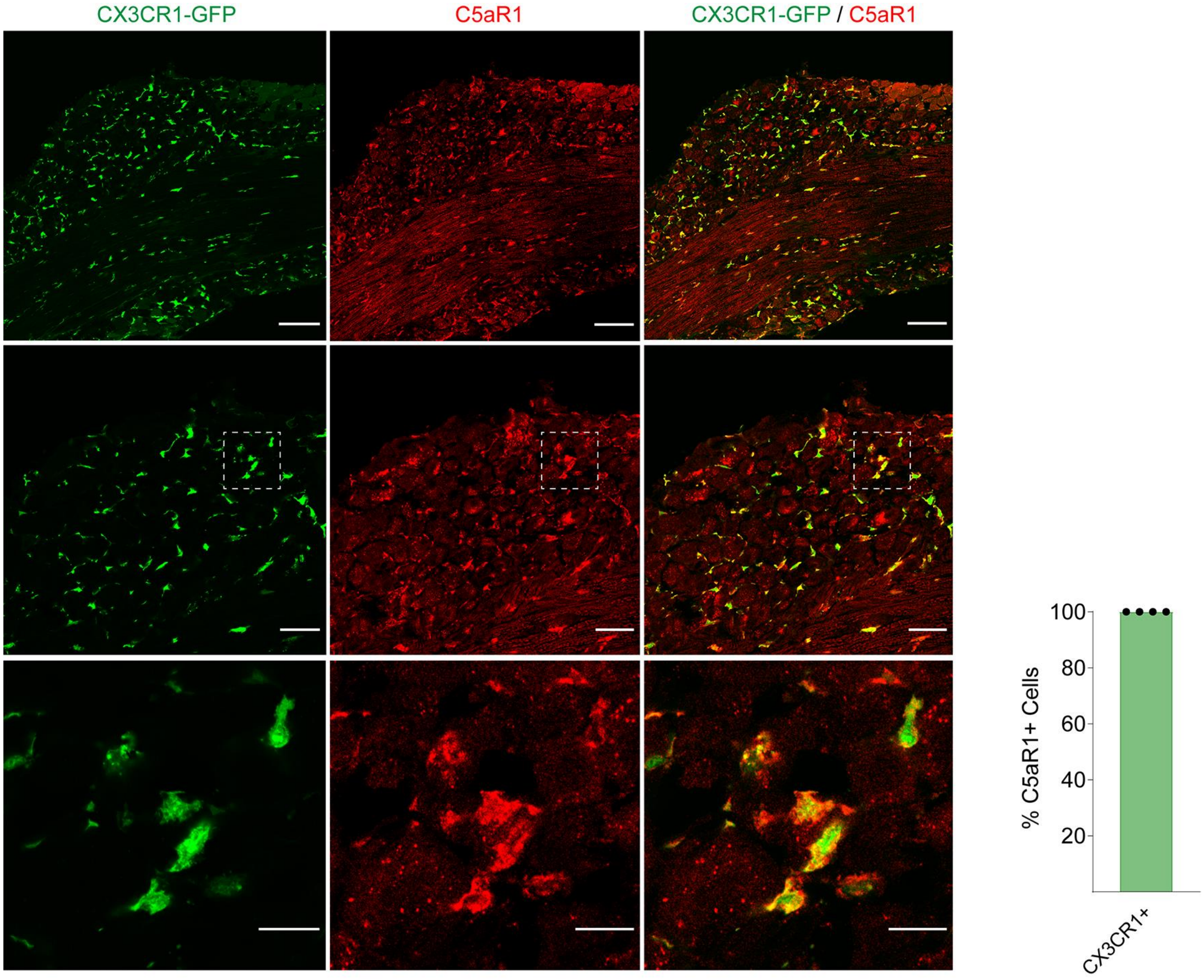


**Figure S5. C5aR1 is expressed in Cx3CR1+ macrophages of the mice DRG.** Representative immunofluorescence and quantification for C5aR1+ cells (red) in CX3CR1+ cells (green, CX3CR1-GFP mice) in the DRG of mice at 7 days after PSNL. Bars in upper panels correspond to 100 µm, the middle to 30 µm and lower panels to 15 µm.

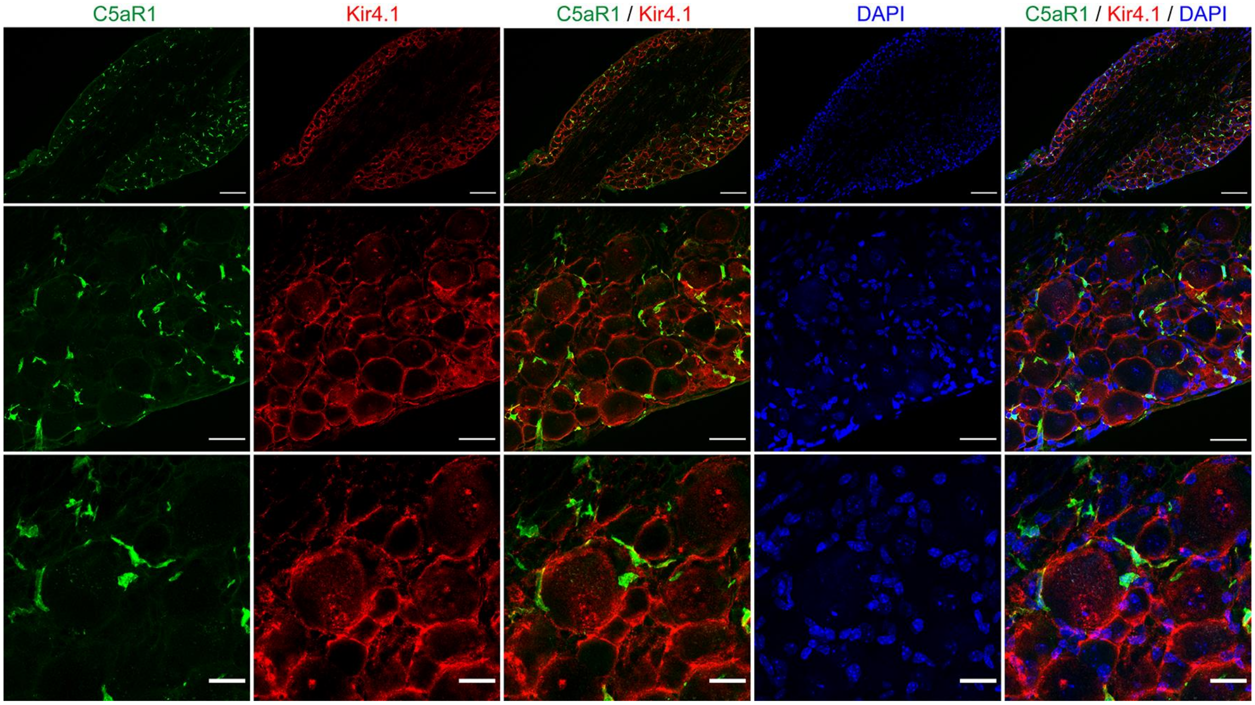


**Figure S6. There is no expression of C5aR1 in satellite glial cells (SGCs) of the DRGs.** Immunofluorescence for C5aR1 (green) and Kir4.1 (red, SGCs) in the DRG samples form of Balb/C WT mice at 3 days after PSNL. Bars in upper panels correspond to 100 µm, the middle to 40 µm and lower panels to 20 µm.

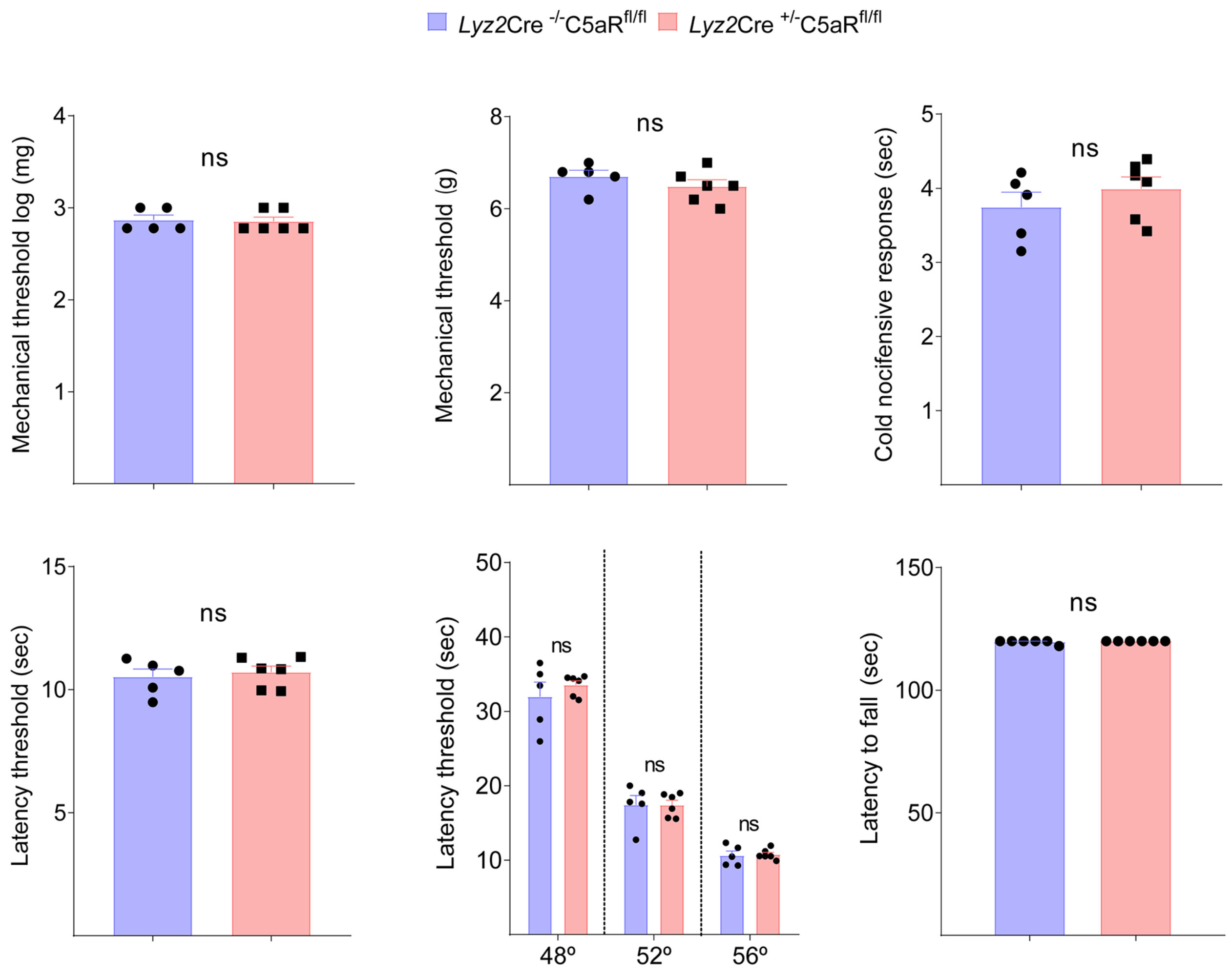


**Figure S7. Conditional deletion of *C5ar1* in *Lyz2*+ cells does not alter nociceptive behaviours.** Physiological nociceptive responses of *Lyz2*Cre$^{+/-}$ - *C5ar*1$^{flox/flox}$ mice compared to littermate controls to mechanical (von Frey filament and electronic von Frey), thermal stimulation (Acetone test, Hargreaves and Hotplate tests) or motor coordination (Rotarod test). Data represented the mean ± s.e.m, *n*=6, and analyzed by non-paired t test - ns, non significant.

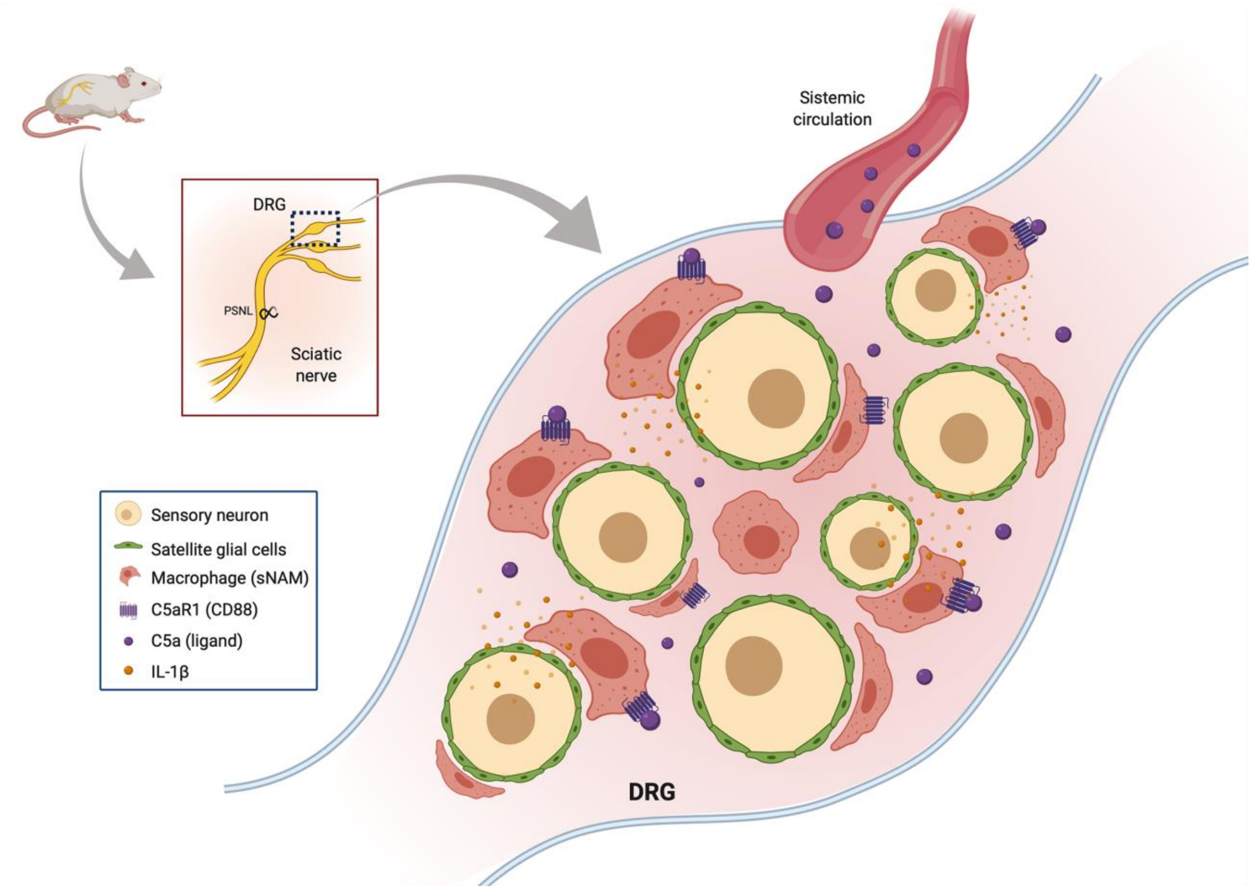


**Figure S8. Schematic representation of the role of C5aR1 signaling in the development of neuropathic pain.** After peripheral nerve injury, C5aR1 signaling in sensory neuron-associated macrophages (sNAMs) of the sensory ganglia drives neuropathic pain through an up-regulation of the pro-nociceptive cytokine IL-1β.

**Supplementary Table 1.** Primers pair used for conventional PCR reactions.

| Target gene | Forward sequence 5’-3’ | Reverse sequence 5’-3’ |
|---|---|---|
| Mouse *C5ar1* | CAAGACGCTCAAAGTGGTGA | TATGATGCTGGGGAGAGACC |
| Mouse *Gapdh* | TGAAGGTCGGTGTGAACGAATT | GCTTTCTCCATGGTGGTGAAG |
| Human *C5ar1* | GCTGCTGAAGAAGCTGGACT | CCCTAACCACGGACTCTTCA |
| Human *Gapdh* | ACCCAGAAGACT GTGGATGG | TTCTAGACGGCAGGTCAGGT |

**Supplementary Table 2.** Primers pair used for RT-PCR reactions.

| Target gene | Forward sequence 5’-3’ | Reverse sequence 5’-3’ |
|---|---|---|
| *Gfap* | AGGGCGAAGAAAACCGCATGACC | TCTAAGGGAGAGCTGGCAGGGCT |
| *Aif1* | TGAGGAGCCATGAGCGAAAG | GCTTCAAGTTTGGACGGCAG |
| *Il1b* | TGACAGTGATGAGAATGACCTGTTC | TTGGAAGCAGCCCTTCATCT |
| *Tnf* | GCCACAAGCAGGAATGAGAAG | AGCAAGCAGCCAACCAGG |
| *Gapdh* | AGGAGCGAGACCCCACTAAC | GTGGTTCACACCCATCACAA |

**Supplementary Table 3.** Antibodies and dyes used for immunofluorescence assays. N/A non-applicable.

| Target | Host | Dilution | Provider | Catalog n. |
|---|---|---|---|---|
| Mouse Tissue | | | | |
| C5aR1/CD88 clone 10/92 | rat | 1:100 | Biorad | MCA2456 |
| Iba1 | rabbit | 1:400 | Wako | 016-20001 |
| GFP (FITC-Conj) | goat | 1:400 | Abcam | ab6662 |
| Nissl (530/615) | N/A | 1:400 | Invitrogen | N21482 |
| Kir4.1 | rabbit | 1:200 | Alomone | APC-035 |
| Ly6G | rat | 1:200 | BD | 551459 |
| Human tissue | | | | |
| C5aR1 | mouse | 1:500 | Biolegend | 344302 |
| Iba1 | goat | 1:500 | Novus | NB100-1028 |
| Nissl (640/660) | N/A | 1:100 | Invitrogen | N21483 |
